# Dental radiography as a low-invasive field technique to estimate age in small rodents, with the mole voles (*Ellobius*) as an example

**DOI:** 10.1101/2024.03.14.585095

**Authors:** Varvara R. Nikonova, Anna E. Naumova, Arman M. Bergaliev, Margarita M. Dymskaya, Anna I. Rudyk, Elena V. Volodina, Antonina V. Smorkatcheva

## Abstract

Most studies which deal with natural populations require a reliable and convenient way of age estimation. However, even rough aging of live individuals is often a real challenge. In this study, we develop a radiographic method of age estimation in *Ellobius talpinus*, a promising model species for population and behavioral ecology. Using a portable X-ray equipment, we radiographed wild, non-sedated animals from the population that had been subjected to extensive mark-recaptures for 3 years. Two molar metrics strongly dependent on age and easy to measure on radiographs were selected: the lengths of synclinal folds of the 1^st^ upper and 1^st^ lower molars. No influence of sex on the molar condition age dynamics was found. Discriminant function analysis based on molar condition and date of radiography in 86 animals of known age classes assigned X-ray images to three age classes (young of the year, yearlings, and 2 years or older) with an accuracy of 99%. Leave-one-out cross-validation yielded 97% of correct assignments. All age estimates for 52 repeatedly radiographed individuals were consistent across images. The analysis of the repeated X-ray images obtained from the same animals showed that 1^st^ lower molars change faster in the first summer of life than later whereas the change rate of the 1^st^ upper molars decreases little throughout life. We propose X-ray technique as a useful alternative to direct skull and dental morphometry for age estimation of wild small mammals, saving the investigator’s time and lives of animals.

## Introduction

Most studies which deal with natural populations require a reliable and convenient method of chronological age estimation. Age-explicit data are essential for conservation biology, population ecology, and the development of life-history theory (Krebs 1999; Holmes and York 2003; Armstrong and Seddon 2008; Zhao et al. 2019; Hecht 2021). For dead mammals, a variety of age indicators can be used including maturity of the dental system, skull or postcranial skeleton, the number of annual layers in the tissues of teeth and bone, and lens mass (Morris 1972; Spinage 1973; Klevezal 2007; Zhao et al. 2019). In some cases, these methods may not be desirable, such as with endangered species or research whose objectives conflict with the removal of individuals (not to mention the ethical aspect). Although examination of tooth replacement and wearing can be executed even on living individuals of some taxa (carnivores - Stander 1997; Landon et al. 1998, ungulates - Gilbert and Stolt 1970, Lipotyphla – Dunaeva 1955; Gonzales-Esteban et al. 2002; bats - Brunet-Rossinni and Wilkinson 2009), for many other mammals such as rodents this is difficult if not impossible. In rodents, incisor width can be a convenient age marker, especially in species with extrabuccal incisors (Kuprina and Smorkatcheva 2019; Caspar et al. 2022), but the reliability of this indicator decreases rapidly with age, and it is useless after the animal stops growing. Estimating age through mark-recapture is time consuming and poorly applicable to species that are difficult to observe and recapture. The most frequently utilized phenotypic characteristics to age live rodents are the mass and length of the body or pelage, but they generally are not sufficiently correlated to age, especially at older age (Ecke and Kinney 1956; Morris 1972; Karels et al. 2004; Lichti et al. 2017; Hulejová Sládkovičová et al. 2019). Some authors determined the age of rodents in vivo by the number of annual layers in the bones of the toe’s phalanges, but these layers are not observed in all species (Klevezal 2007). During the last decades, various molecular age markers, such as DNA methylation, racemization, pentosidine and telomere length, have been proposed (see Zhao et al. 2019 for review). The accuracy of these methods varies and is sometimes quite high, but they are time-consuming and expensive. In addition, they require remote laboratory analyses and, therefore, are not suitable when it is desirable to estimate the age of an animal immediately after its capture. Behavioral markers of age as e. g. parameters of distant calls in the cheetah *Acinonyx jubatus* (Klenova et al. 2023) have been described for some carnivores, but revealing of similar age indicators in rodents is unlikely. Thus, non-invasive or low-invasive aging of rodents is often a real challenge (Klevezal 2007; Zhao et al. 2019).

With technological advancements of recent years, skeletal, cranial and teeth features of live animals can be examined using radiography without killing animals, even during brief capture. Dental radiography is a simple method that allows traditional age indicators such as molar condition to be applied. The possibility of recording age-related dental changes on radiographs has been already shown for the Martino’s vole, *Dinaromys bogdanovi* (Kryštufek et al. 2000) and bank vole, *Myodes glareolus* (Alibhai 1980). Whereas these authors obtained radiographs from dead animals, we aimed at the development of a radiographic method of age estimation in live rodents, northern mole voles (*Ellobius talpinus*).

Subterranean lifestyle, cooperative breeding and unusual life history traits (Letitskaya 1984, Evdokimov 2001, Moshkin et al. 2007; Kaya and Coşkun 2015, Smorkatcheva et al. 2016; Smorkatcheva and Kuprina 2018) make mole voles excellent experimental models for population and behavioral ecology. The age structure of *E. talpinus* populations is complex due to extremely long (by arvicoline standards) lifespan which is up to six years even in free-ranging animals (Evdokimov 2001). Although abundant throughout most of its range, this species is considered endangered in Ukraine (Akimov 2009). Clearly, establishing a non-invasive method of age determination is critically important for many studies using mole voles as well as monitoring this species. Very young mole voles, up to the age of approximately 2 months, can be readily recognized by their small size, grayish (in the case of the brown morph) pelage, and narrow upper incisors. Incisors’ width can be used for rough aging of mole voles younger than 3-5 months (Kuprina and Smorkatcheva 2019). The fact that molars in *Ellobius* develop roots during postnatal growth offers an opportunity to use teeth structure as an age indicator. Evdokimov (1997) developed a method for dividing northern mole voles into age groups based on the length of tooth roots, and Kropacheva and coauthors (2018) provided the equation for the relationship between the length of tooth roots and the age of laboratory-born northern mole voles. However, no non-destructive method that would allow even rough aging of adult mole voles is currently available. In this work, we took the advantage of recent progress in portal digital radiography to develop such a method.

Ideally, the description of age-related changes should be based on the study of individuals whose accurate age is known, which is only possible when working with laboratory-born animals. We had a laboratory colony of a closely related species, *E. tancrei*, at our disposal. We used these animals to develop the method of X-raying live mole voles and to preliminary reveal age-dependent parameters that can be measured on radiographs. Subsequently, when comparing age-related dental changes in laboratory *E. tancrei* and wild *E. talpinus*, a significant discrepancy was found, which most likely reflected the influence of conditions rather than species differences (Lowe 1971 and Abe 1976 for *Myodes*; Kropacheva et al. 2018 for *E. talpinus*). Therefore, this study is entirely based on the data from free-ranging animals.

## Material and Methods

### Study species and population

The study site is located in the Saratov Region (Russia), four kilometers west from Dyakovka village (50.71°N, 46°71′E). This territory lies on the border of steppe and semi-desert zones. The vegetation cover is represented by psammophytic-steppe and meadow-steppe types of plant communities. As part of the projects to study the population genetic structure, behavioral ecology, and acoustic communication of the northern mole vole, mark - recapture method has been used on an area of about 25 ha since spring 2021, with 3-4 trapping sessions per year: May 17-24, July 16-19 and August 24 - September 5 2021, May 16-28, July 16-August 9, and September 26 - 30 2022, May 2 -13, June 21 - July 3, August 8 - 22, and September 27 - October 6 2023. Population density of mole voles, estimated as the number of animals known to be alive, has been consistently high, reaching in some places 20 individuals per hectare. In the Saratov Region, the reproduction of *E. talpinus* is seasonal. We have observed juveniles up to the beginning of September and lactation females up to the middle of August. There is no exact data on the beginning of the breeding season but most probably it falls at the beginning of March. *E. talpinus* is a highly social species living in extended family groups with a complex kinship and age structure. Within each family, only one female typically reproduces (Evdokimov 2001; our observation). According to the published data for Chelyabinsk Region, Russia (Evdokimov 2001), which appears to be consistent with our observations for the studied population, each female breeder delivers 1 or 2 litters per season. The minimum interbirth interval, approximately equal to the duration of pregnancy, is about 30 days (Zadubrovskaya et al. 2020).

### Trapping and X-ray imaging

The animals were captured with metallic-spiral live-traps (Golov 1954 with modifications) placed into burrow tunnels. At first capture, each animal was tagged with microtransponders 1.25*7 mm (Star Security Technologies Co., Shanghai, China) for further identification. For each individual, the capture date, accurate GPS coordinates of the capture site (to the nearest 0.0001°), sex, pelage condition (grayish, brown or molting), body weight (to the nearest 0.1 gram), and joint width of both upper incisors (measured with electron caliper to the nearest 0.01 mm) were recorded. In addition, bioacoustical data were collected (Dymskaya et al. 2024), and distal phalanges of one or two toes were clipped to be used for the genetic analyses (Rudyk et al., in prep.).

Since July 2022, obtaining radiographs has been added to the described procedures. We performed X-rays without sedation of mole voles in order to prevent side effects that can result in death (Hawkins 2020), especially given the stress experienced by captured animals and hot ambient temperatures. This work is a part of a long-term population study, which does not allow the removal of any individuals from the population. Therefore, the tradeoff between the high quality of radiographs and the safety of animals was resolved in favor of the latter. The required positioning of the animal’s head was achieved by placing the animal into a falcon tube (50 ml) with the end of the bottom cut off (Fig. SI1 and SI2, Supplementary Information). Wright lateral radiographs were taken from an animal’s cranium with a portable X-ray equipment (Rexstar LCD, Korea) and Dental Radiovisiography Sensor EzSensor 1.5 (Vatech). The exposure time was 0.18 s; the distance between the end of the cone and the tube with an animal was 15 cm. The resulting images were stored and processed in the EzDent i. 3 software (http://www.ewoosoft.com/). We usually took 2 - 5 images from each animal to ensure the obtaining of radiographs of acceptable quality and selected the best one for the analysis.

Each animal was released exactly into the hole where it had been caught. Each individual was subjected to this procedure once within one trapping session, but if recaptured in subsequent sessions, it was radiographed again.

### Age classes and known-age mole voles

Given the available data on the development of captive mole voles (Kuprina and Smorkacheva 2019 for a sibling species, *E. tancrei*) and reproductive seasonality, the animals with incisor widths less than 3.15 mm could be reliably identified as young of the year. Accordingly, the radiographs of most animals (hereafter «*known-class images”*) were assigned to one of three age classes (Table. 1): class 1 - images from young of the year; class 2 - images from the animals that were captured as young last year, i. e. yearling; class 3 - images from the animals known to survive at least two winters. We also had radiographs from mole voles known to survive *at least one winter* (hereafter referred to as “*unknown overwintered images*”).

Thirty mole voles could be aged to the nearest 5-10 days based on their appearance at first capture. The smallest mole voles to trap had body mass of 18-25 g, short gray pelage, and incisors width up to 2.60 mm; they were caught very rarely and only after all the older members of their family had been trapped. Given that juvenile northern mole voles open their eyes at 22-23 days with a body mass of 15-20 g (Letitskaya 1984; Smorkatcheva, unpublished), these individuals were approximately 30 ± 5 days old. The youngest animals that readily entered traps were also grayish, but obviously larger and typically heavier, with body mass up to 40 g and incisors width up to 3 mm. Their approximate age could be estimated as 40 ± 10 days. We realize that these age estimates may not be accurate and are actually relative, reflecting stages of development rather than absolute age. However, due to the absence of more reliable information, we used mean values of these age ranges to evaluate the predictability of yearlings’ absolute or relative age based on dental characteristics. We calculated the approximate age of the known-age mole voles at the time of radiography by adding their presumed age at first capture and the interval from first capture to the radiography. All radiographs of these individuals are referred as *“known-age images”*.

### Analysis of radiographs and selection of age-dependent characters

A total of 245 good quality radiographs were selected for this study (132 male radiographs from 91 individuals and 113 female radiographs from 78 individuals). These included 144 known-class images obtained from 86 animals (Table 1) and 20 unknown-class overwintered images from 16 animals. The remaining 81 images belonged to 71 animals whose age class at the time of X-ray is unknown; they also were used in some analyses (see Statistical analysis). Fifty-two animals were X-rayed several times, in different trapping session or in different years. Therefore, for some animals we had images belonging to different classes (unknown overwintered and class 3, unknown and unknown overwintered, class 1 and class 2, or class 2 and class 3, Table 1). Among the known-class radiographs, 49 were known-age images from 30 animals.

**Table 1.** Sample sizes of known-class radiographs used in this study.

|  | Age class | males | females | In total |
| --- | --- | --- | --- | --- |
| Number of animals | Only 1 | 31 | 30 | 61 |
|  | Only 2 | 5 | 2 | 7 |
|  | 1 and 2 | 3 | 2 | 5 |
|  | Only 3 | 6 | 4 | 10 |
|  | 2 and 3 | 2 | 1 | 3 |
|  | **In total** | **47** | **39** | **86** |
| Number of radiographs | 1 | 50 | 47 | 97 |
|  | 2 | 20 | 7 | 27 |
|  | 3 | 11 | 9 | 20 |
|  | **In total** | **81** | **63** | **144** |

The selected radiographs were processed through the EzDent i. 3 software (http://www.ewoosoft.com/) which allows image rotation, scaling, and measuring length, angles and density.

We screened the radiographs obtained from the known-class animals and those from the same animals in different trapping sessions to reveal age-dependent characters. A number of candidate characters (length of molar roots and molar crown, the maximum length of the grinding surface, angles of inclination of molars, and several other metrics and indexes, as well as the density of some skull structures) were then evaluated for interobserver and intraobserver variability. Measurements were performed with a digital tool (accuracy 0.01 mm) by two observers (VRN and AEN) blind to the animal’s age class. As a result, we selected two parameters that seemed to change with age and were least prone to measurement errors: the length of the second synclinal fold of the first upper molar (USF) and the length of the second synclinal fold of the first lower molar (LSF, Fig. 1). For these metrics, the average interobserver difference (the difference between measurements by two observers divided by the average value) was 0.06 and 0.12, respectively. Intraobserver differences (the difference between two measurements by the same observer divided by the average value) were 0.04 and 0.07 (VRN) and 0.06 and 0.09 (AEN) for USF and LSF, respectively. Given a relatively low precision of measurements, the mean of all individual measures by two observers was used.

**Fig. 1.**
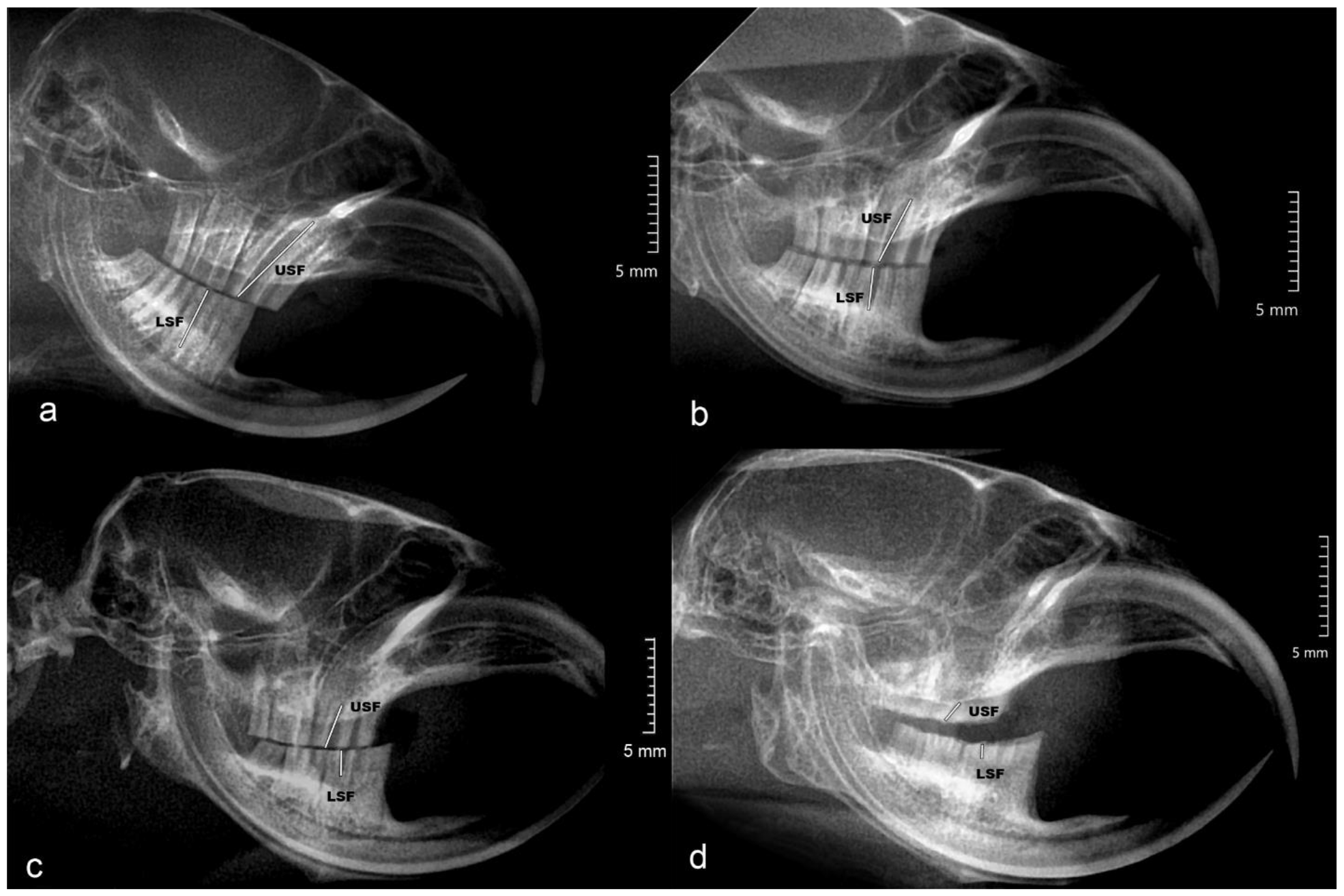
Skull radiographs of mole voles from three age classes: a - class 1 (young of the year), b - class 2 (yearling); c, d - class 3 (survived two or more winters). USF – synclinal fold of the 1^st^ upper molar; LSF – synclinal fold of the 1^st^ lower molar

### Statistical analyses

First, we tested the appropriateness of the selected molar metrics for discriminating between age classes using all known-class radiographs (n = 144). We examined, separately for two sexes, the basic USF and LSF statistics for each age class to test whether some or both characters can be directly used to distinguish between the three age classes. Then we estimated the correlation between USF and LSF with Pearson’s correlation coefficient, using all available radiographs (n = 245) and combining data for males and females. Next, a principal components analysis (PCA) was performed (again, using all 245 radiographs) to eliminate redundancy due to the high intercorrelation between two metrics by summarizing them into a single factor, molar condition.

We used a linear mixed model analysis (LMM, *lme4* package in R 4.3.1) to investigate the effects of age class, sex and Julian day of radiography as well as their interactions (fixed effect predictors) on molar condition. Animal’s identity was fitted as a random term. All known-class radiographs were included in this analysis (n=144). We compared models based on maximum likelihood estimates. Akaike’s Information Criteria corrected for small sample size (AICc) was used to guide model selection (Mazerolle 2023). In cases of similar support (ΔAICc < 2), the model was preferred due to a lower number of parameters. The appropriateness of the best model was checked via diagnostic of observations influence through the Cook’s distances (Cook 1979) (Di < 1) and via analysis of residuals using the Shapiro-Wilcoxon normality (Royston 1995) (p > 0.05) and the Breusch-Pagan homoscedasticity (Krämer and Sonnberger 1989) (p > 0.05) tests. Based on the results of the LMM, we conducted all the following analyses on the pooled data for males and females.

We performed the discriminant function analysis (DFA, Statistica 12) to test how well molar condition can be used to classify 86 known-class images (one image per an animal, class 1: n=61; class 2: n=13; class 3: n=12) into age classes. For those individuals X-rayed in both years, the latest radiograph was used to increase the sample size for older age classes. When we had several radiographs per individual per year, the best quality image was selected. Julian day of radiography was included as a second predictor. Kolmogorov–Smirnov test was used to check multivariate normality of data distribution. Prior probabilities were based on group size. The test error was estimated using leave-one-out cross-validation (*caret* package in R 4.3.1).

We applied the obtained classification functions to classify all remaining radiographs (i. e. 20 unknown-class overwintered images, 81 unknown-class images and 58 known-class images that were not used for classification functions computation). This was done in order to test whether the yielded classification of different radiographs from the same animals would contain any contradictions, and whether any radiographs from unknown overwintered animals would be misclassified as young of the year (age class 1).

We used a two-way ANOVA (Statistica 12) to compare the age-related dynamics of USF and LSF for those known-class individuals that were X-rayed during several trapping sessions. In this analysis, a metric (USF/LSF) was a within-group factor, and interval class was a between-group factor with three levels: intervals between two radiographs taken in the first summer of life (n = 23), intervals between two radiographs taken in the first and second summers of life (n = 6) and interval between two radiographs taken in the 2nd and 3rd summers of life or later (n = 4). The rate of change was determined as (measurement 1 - measurement 2)/interval between surveys (in months). The obtained values were log-transformed to improve normality of data distribution. The significance of pairwise differences was tested with the Unequal N HSD-test.

Finally, we estimated the predictability of age (judging by the R^2^ values and prediction intervals on scatter plots) based on USF and LSF, using the radiographs from known-age mole voles (n=30). For each known-age animal, the last or only image taken in the first summer of life was included in the analysis. The resulting range of ages covered a period of life approximately from 1.5 to six months.

In all analyses significance levels were set at 0.05, and two-tailed probability values are reported.

## Results

The comparison of basic statistics for USF and LSF between age classes suggested the usefulness of these metrics for age estimation: they significantly decreased from age class 1 to age class 3 in both sexes (Fig. 2). In our samples, there was no overlap between yearlings and the two older age classes in USF. However, there was some overlap in both parameters between age classes 2 and 3, and therefore neither was sufficient to correctly assign animals to the classes (Fig. 2).

**Fig. 2.**
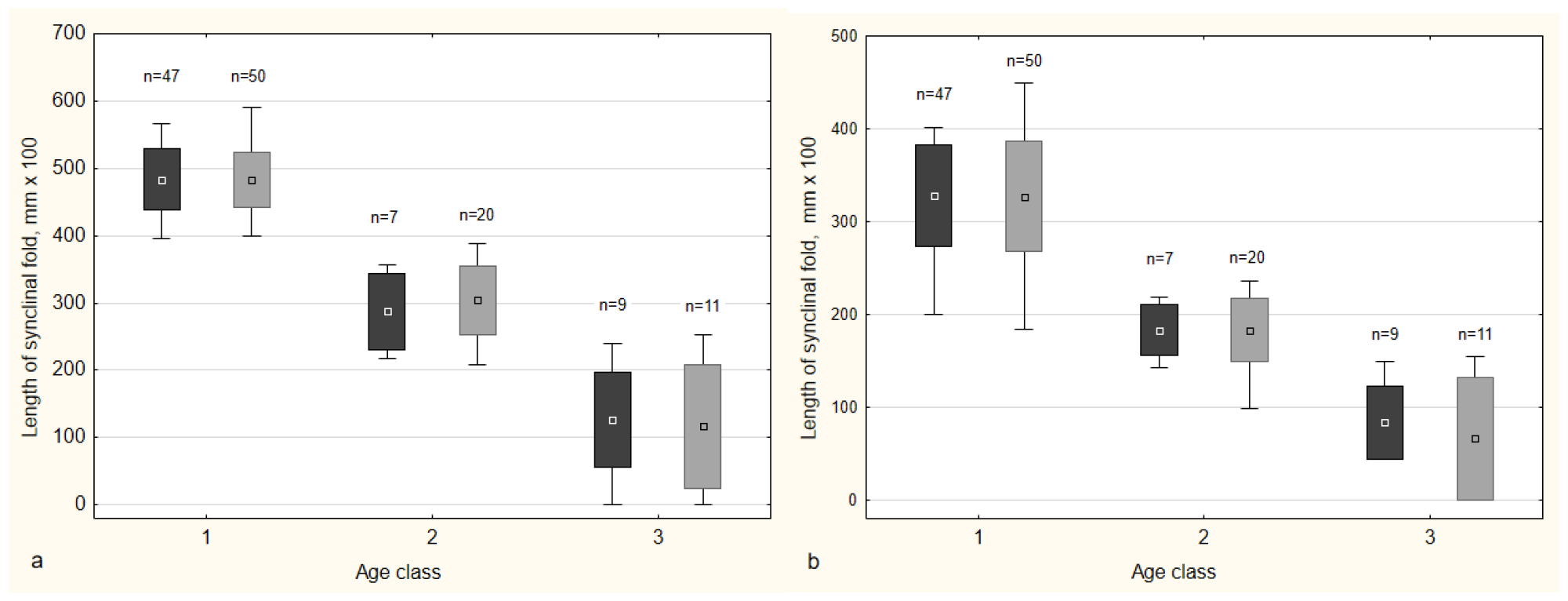
Variation in synclinal fold lengths of (a) upper and (b) lower first molars in *E. talpinus* from three age classes (mean ± SD and limits). Dark gray boxes - females, light gray boxes - males

**Fig. 3.**
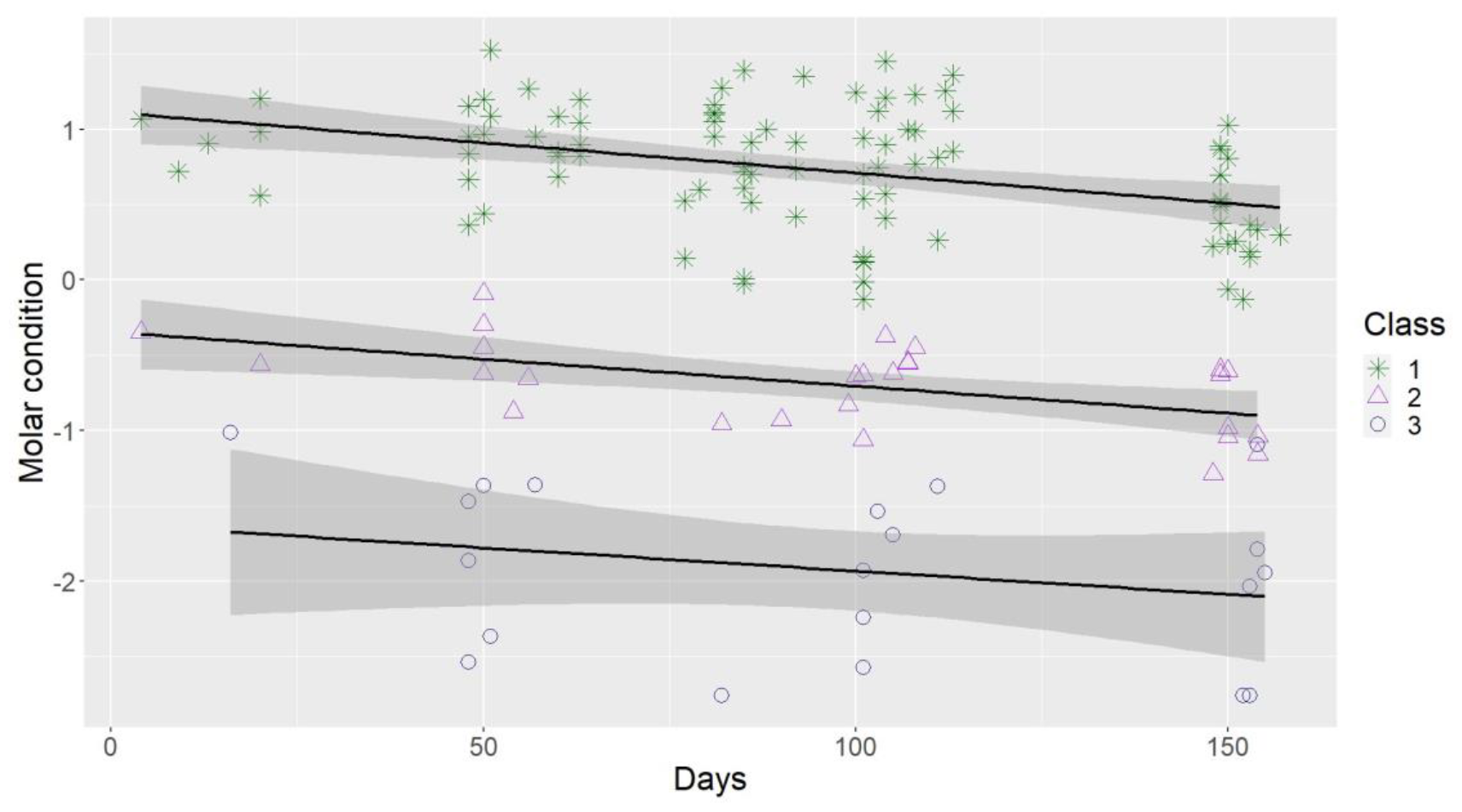
Relationship between Julian day of radiography and molar condition in mole voles of three age classes based on 144 radiographs from 86 individuals

There was a strong correlation between USF and LSF (r = 0.92; p < 0.001). Principal component analysis yielded PC1 which accounted for 96% of the total variation (factor loading 0.98). In the following, we use the PC1 as an indicator of molar condition.

In the LMM analysis, age class, Julian day, and the interaction of day and age class were the best combination of explanatory factors associated with molar condition (Table 2). The model which included the same predictors plus sex had similar support but a larger number of parameters. The best model explained 97% of variation in molar condition. The parameters of the top prediction regression model for molar condition are presented in Table 3. Molar condition decreased from spring to autumn in animals of all age classes, somewhat faster in yearlings (Fig.3).

**Table 2.** Model selection results from the mixed-model regression analysis of molar condition in mole voles based on 144 radiographs from 86 individuals. All models included a random effect of animal’s identity. ΔAICc - differences in AICc values relative to the best model. Lowest AICc = 79.85

| Fixed effects in model | df | $\Delta\text{AICc}$ |
| --- | --- | --- |
| Age+day+age*day | 8 | 0.000 |
| Age+day+age*day+sex | 9 | 1.850 |
| Age+sex+age*sex+day+age*day | 11 | 5.108 |
| Age+day | 6 | 13.917 |
| Age+sex+day | 7 | 15.670 |
| Age+sex+age*sex+day | 9 | 20.178 |
| Age | 5 | 56.156 |
| Age+sex | 6 | 58.266 |
| age+sex+age*sex | 8 | 62.645 |

**Table 3.** Parameters of the best linear mixed model of molar condition in mole voles based on 144 radiographs from 86 individuals.

| Parameter | Estimate (Std.Error) | *t* |
| --- | --- | --- |
| Intercept | 0.598 (0.044) | 13.723 |
| Age class 2 | -1.419 (0.062) | -22.962 |
| Age class 3 | -2.268 (0.092) | -24.693 |
| Day (standardized) | -0.256 (0.022) | -11.837 |
| Age class 2 : day (standardized) | 0.174 (0.040) | 4.308 |
| Age class 3 : day (standardized) | 0.200 (0.052) | 3.839 |

DFA based on 86 known-class radiographs confirmed that molar condition and day of radiography, taken together, ensured the discrimination between age classes (Wilks’ lambda = 0.15; χ2=158.9; df = 4; p<0.0001). The model explained 99% of the variation in the variable.

The plot of the canonical scores for the first two discriminant functions illustrates the separation among age classes (Fig. 4). The first discriminant function was highly correlated (*r* = 0.95) with the molar condition, whereas the second discriminant function was highly correlated with day of radiography (*r* = 0.99). The first discriminant function accounted for 99.9% of the grouping variation. It was weighted most heavily by the molar condition (standardized coefficients -1.05 and -0.31, respectively, respectively).

**Fig. 4.**
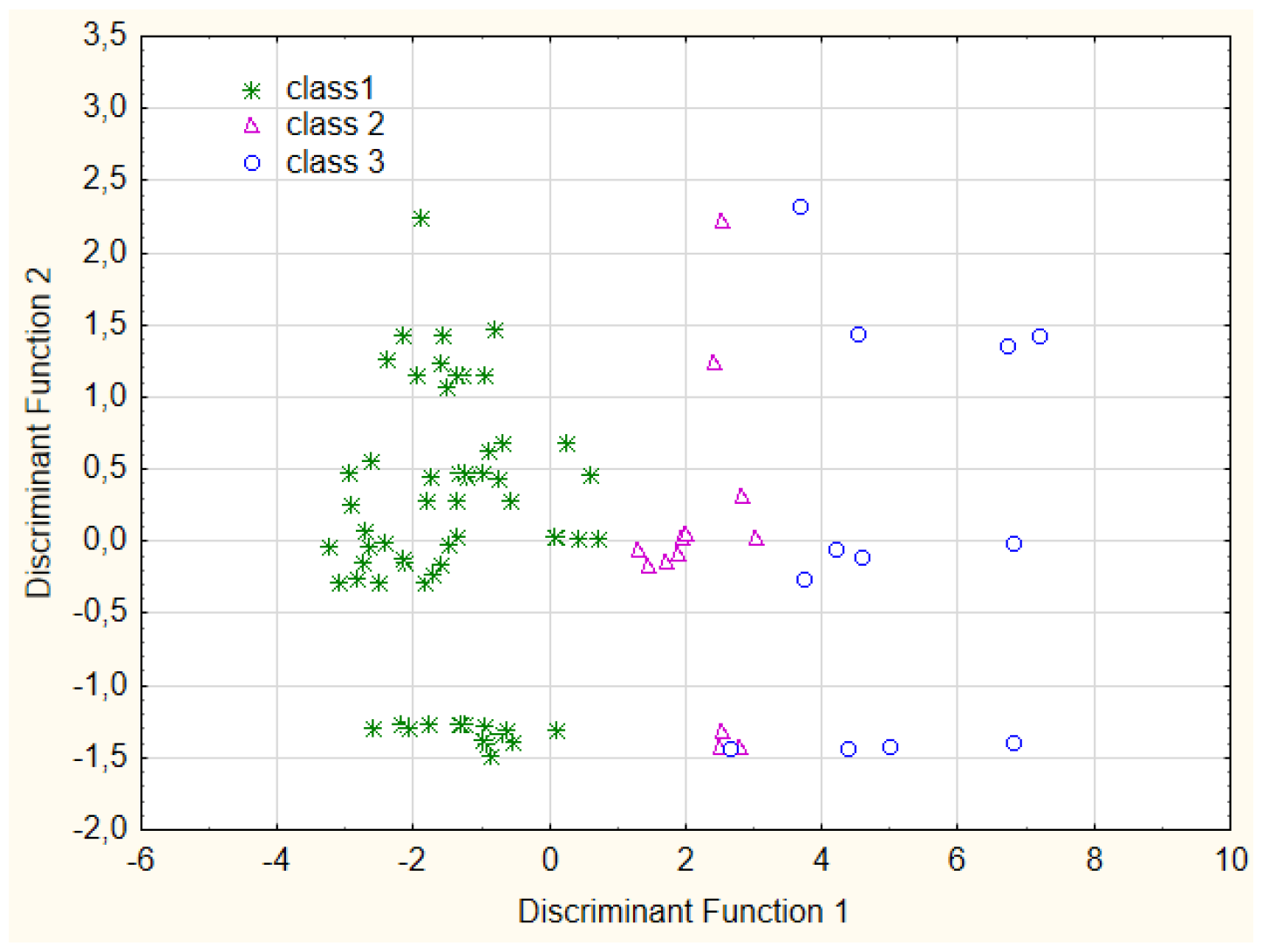
Scatterplot illustrating DFA results for classifying 86 images from 86 known-class mole voles

The following discriminant equations were obtained:

Class 1: -25.28 + 9.05 molar condition + 19.28 day

Class 2: -20.43 + 0.41 molar condition + 0.17 day

Class 3: -23.63 -6.82 molar condition + 0.14 day

The predictive accuracy of the model for the analysis sample was 0.99. Only one radiograph from class 3 was mistakenly assigned to class 2 whereas all images from classes 1 and 2 were classified correctly. The predictive accuracy of the cross-validation sample was 0.97 (95% CI: 0.901 - 0.993).

All 20 radiographs of the unknown overwintered animals were assigned by DFA to either class 2 or class 3. All age estimates for repeatedly radiographed individuals (2 to 5 images from each of 52 animals) were consistent across images: the images from the same individual were always assigned to the same class if they had been obtained in the same year and to different classes if they had been obtained in different years.

Two-way ANOVA revealed the difference in age-related patterns of USF and LSF. Both main effects, as well as their interaction, were highly significant (interval class: F2,30 = 12.6; p = 0.001; metric: F1,30 = 8,9; p = 0.006; interval class X metric: F2,30 = 13,6; p < 0.001). LSF decreased faster during the first summer than later in life, and LSF decreased faster than USF early in life. The decreasing rate of USF, in contrast to LSF, changed little throughout life (Fig. 5).

**Fig. 5.**
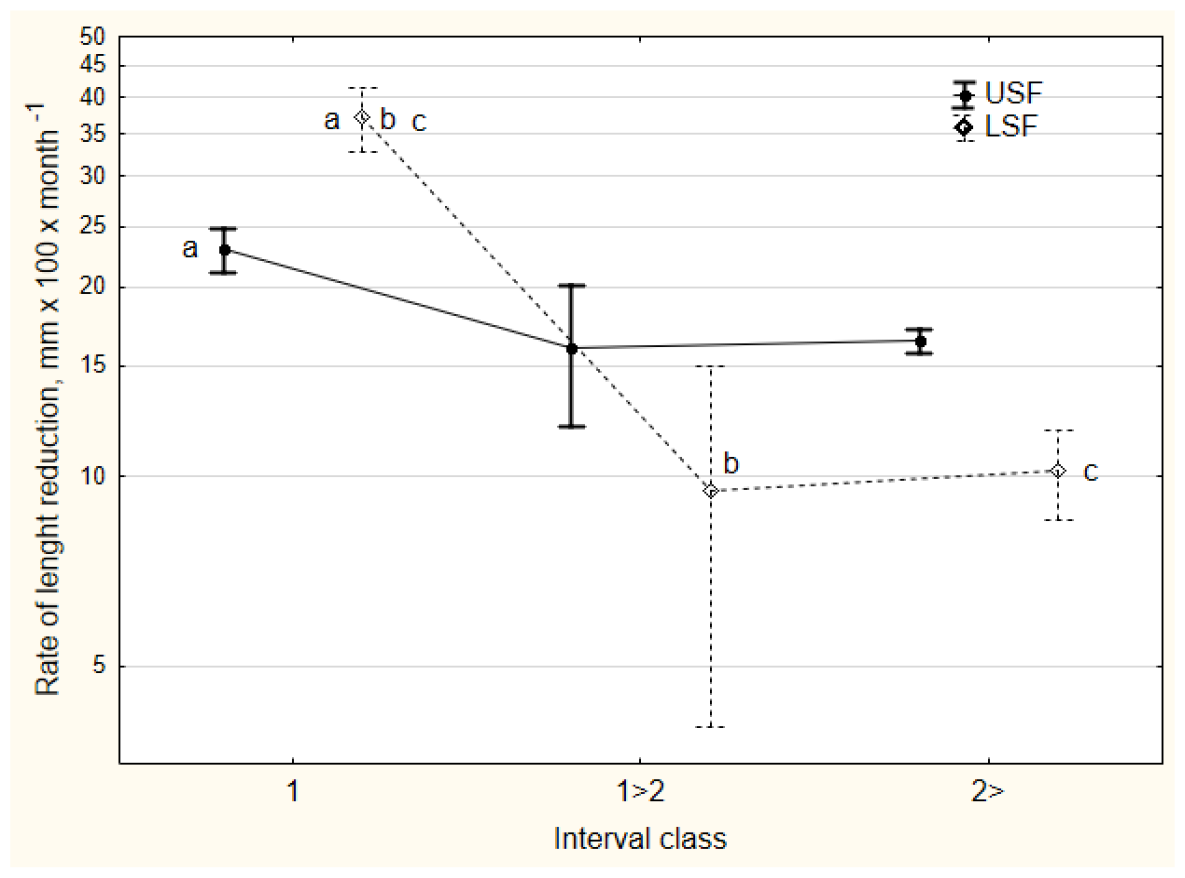
Rates of decreasing in synclinal fold lengths of lower (LSF) and upper (USF) 1st molars (mean±*SE*) during different periods of life. Interval classes: 1 - between two radiographs taken in the first summer of life; 1>2 - between two radiographs taken in the first and second summers of life; 2> - between two radiographs taken in the 2nd and 3rd summers of life or later. (a: p < 0.05; b and c: p < 0.001).

In 30 known-age young of this year, the fit between age and LSF was better than fit between age and USF although the determination coefficients differed non-significantly (p = 0.058). The predictability of age based on each of two metrics was too poor to be used, for example, to distinguish cohorts of yearlings (Fig.6).

**Fig. 6.**
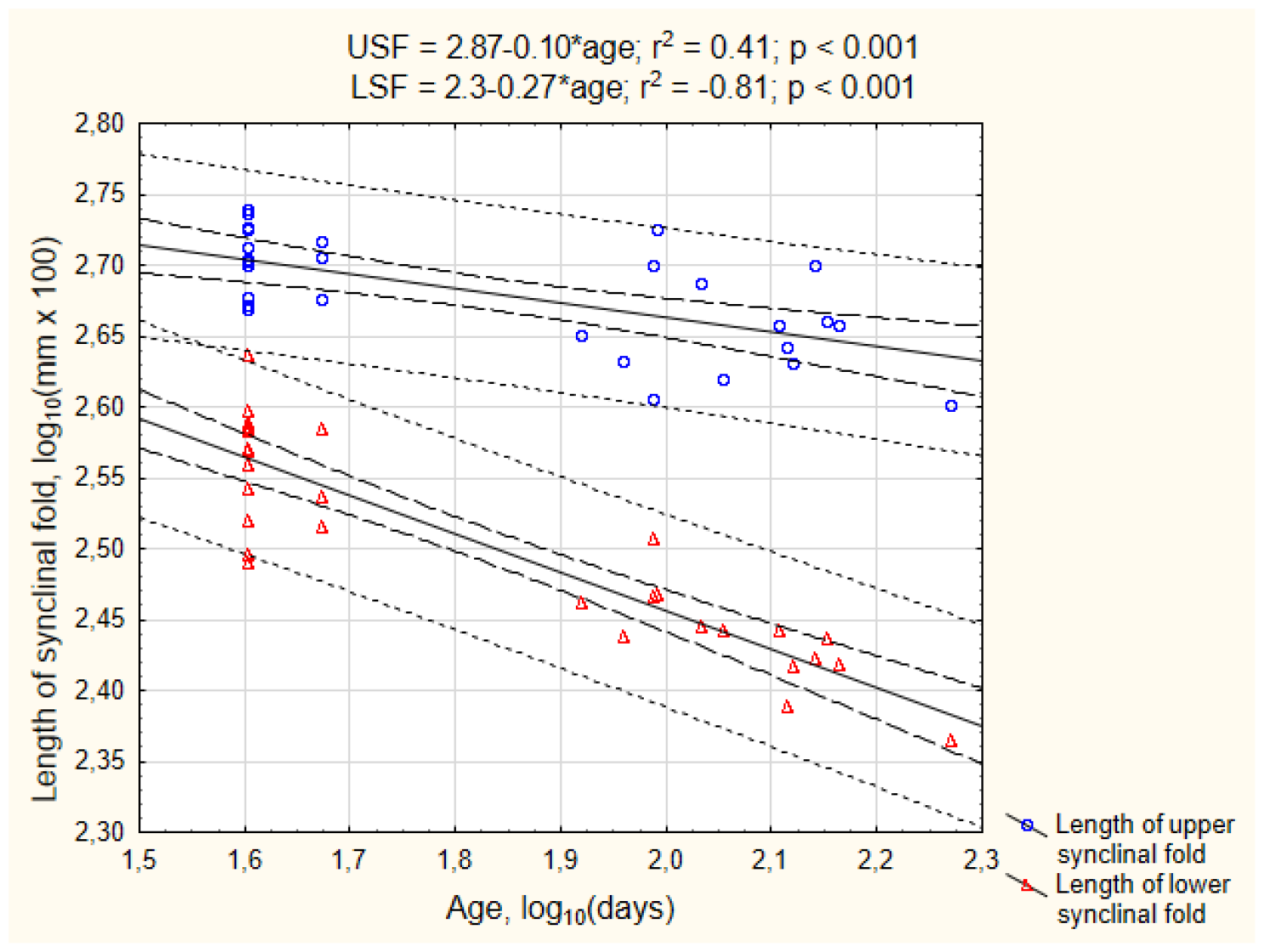
Scatterplot of the length of the synclinal fold of upper (USF) and lower (LSF) 1st molars against age (in days) for 30 known-age mole voles (solid line - log-linear regression model, dashed line - 95% confidence intervals; dotted line - 95% prediction interval)

## Discussion

In this work, we applied dental radiography for estimating the age of the northern mole vole, which is a valuable model object for ecological and behavioral research due to the subterranean specialization and sociality (Evdokimov 2001, Moshkin et al. 2007). Molar roots length is traditionally used as age indicator for rhizodont voles, including *Ellobius* (Tupikova et al. 1968; Lowe 1971; Viitala 1971; Abe 1976; Alibhai 1980; Gustafsson et al. 1982; Kryštufek et al. 2000; Evdokimov 2001; Kropacheva et al. 2018). According to Evdokimov (1997, 2001), the 1st lower molar in *E. talpinus* starts developing roots during the first summer of life. However, even in the radiographs obtained from old animals, molar roots were not always clearly visible (Fig.1 c, d), and only in some radiographs were they measurable. Therefore, two other molar metrics were selected. One of them, the upper synclinal fold of the 1^st^ molar (USF), is apparently sufficient to separate young of the year from overwintered individuals. For further age differentiation of overwintered animals another parameter, the 1^st^ lower molar synclinal fold (LSF), and the date of X-ray should also be taken into account. Our model allows us to discriminate between three age classes with much greater reliability than the previously described method based on the direct dental measurements of only the 1st molar roots (Evdokimov 1997). The results of the LMM analysis suggest that the rude categorization of *E. talpinus* to age classes can be done without taking sex into account, although, in view of the paucity of the data for the adult groups, the effect of sex on molar condition changes cannot be ruled out.

Based on the lifespan reported for wild northern mole voles (Evdokimov 2001), class 3 may include animals 3-6 years old. The heterogeneity of this class is confirmed by high intra-class variability of USF and LSF (Fig. 2), as well as the diversity in molar root condition (Fig. 1 c, d). The scatterplot of the canonical scores (Fig. 4) also hints that class 3 may combine animals from at least two discrete subclasses. Within the timeframe of our mark-recapture study, it was not possible to obtain the data necessary to statistically test this assumption. We hope that in the future we will elaborate our model to differentiate between oldest age classes. However, the inability to discriminate between old and very old animals may not be critical for many researchers because the latter category usually constitutes only a small proportion of a population. In *E. talpinus* from South Ural, a pooled group of 4-6 years old mole voles constitutes only 6.5% of the population (Evdokimov 2001).

The breeding season of mole voles in the studied population is extended over four months, so class 1 encompass several cohorts of young animals. LSF overperforms USF as a potential age indicator for young of the year. To create a model for more accurate age estimation within class 1, it is necessary to know exact dates of birth, but obtaining such information would require special research. However, attempts to achieve greater accuracy in absolute age determination based on molar condition do not seem particularly promising, given the very likely inter-cohort differences in root development and teeth attrition (Lowe 1971; Olenev 1989). In the case of the focal population of *E. talpinus*, the relative age of young of the year can be estimated based on their pelage condition and incisors width (Kuprina and Smorkatcheva 2019) for most of the trapping period.

Although X-rays have been widely used for quite a long time to determine the age of mammals, such studies focus mainly on large, rare or long-lived species (carnivores - Jenks,1984; Dix, Strickland, 1986; Zeiler 1988; Nicholson et al. 2020; cetaceans - Read et al. 2018; Barratclough et al. 2019; ungulates - Flinn et al. 2013). To our knowledge, this study is the first attempt to apply radiography for age determination in live wild rodents. Most aging methods developed on laboratory-born animals require validation before use in the field; without such an additional study, at best they can be used to estimate relative rather than absolute age. This is especially true for dental indicators because growth rates of molar roots and attrition rate depend on the diet and vary during the year (Lowe 1971; Abe 1976; Olenev 1989; Evdokimov 2001; Kropacheva et al. 2018; Kropacheva et al. 2021). We believe that our method is applicable to any mole vole population living in areas with similar seasonality and soil and vegetation characteristics. In regions with very different conditions, adjustments may be necessary.

The main disadvantages of using animals from a natural population for developing age estimation methods were already mentioned: (i) not knowing the age of some old individuals; (ii) not knowing the exact age of any individual; and (iii) a biased sampling with a large predominance of younger categories. In the case of mole voles as well as other social subterranean rodents, field studies of the relationship between age and morphological traits may be potentially confounded by the effect of social/reproductive status (although for at least one species a specially conducted study did not find such an effect – Caspar et al. 2021). However, if critically necessary, these problems can be overcome by the extension of a field study, intensive trapping at the very beginning of the reproductive period, and monitoring of the reproductive condition of tagged individuals.

To summarize, the dental X-ray technique has proved a useful alternative to direct dental/skull morphometry for age estimation of wild small mammals, saving the investigator’s time and lives of animals. As a tool for age assessment, radiography may be especially in demand for monitoring rare species of rodents (dormice, jerboas, and many representatives of Myomorpha) or other taxa and for demographic studies of species with long life-histories. The way of obtaining radiographs used here, namely X-raying non-sedated animals, was dictated by the objectives of the main study of which this work is a part. Because it was difficult to manage complete immobility, perfect symmetry and overlap of the teeth, the quality of our images was not ideal, but it was sufficient to achieve our goal. It is possible that chemical immobilization may be required or desirable for X-ray in other species and for other purposes. The full immobilization could possibly improve the quality of X-rays and, as a result, reduce the error in age determination. Moreover, immobilization of an animal during radiography would allow the use of age indicators other than teeth or skull characters. The degree of fusion of the epiphyseal plates and pelvis features, unlike the indicators dependent on tooth wear, may be appropriate for age estimation in arhizodont species of rodents (Tarasov 1966; Zudova et al. 2017).

## Supporting information

Skull radiography of live, non-sedated mole voles

## Acknowledgments

We are grateful to N. V. Kryukova for her valuable advice on X-ray methods. Many thanks to I. A. Volodin, A. A. Panyutina, A. N. Kuznetsov, M. S. Gribanova, A.O. Fedosov, M. Y. Vdovina, P. S. Cherepenko and E. D. Kolosova for their participation in our long-term field study. We thank A.V. Tchabovsky for his help in the organization of field work. Early reviews of this manuscript provided by I. A. Volodin significantly improved its quality.

## Ethical notes

All procedures involving animals were in compliance with the ASAB/ABS Guidelines for the Treatment of Animals in Behavioural Research and Teaching (Buchanan et al. 2012) and to the national laws of Russian Federation. The experimental protocols used in this study were approved by the Specialized Ethics Committee for Animal Research of the St. Petersburg State University (№№ 131-03-2 from 02.02.2021 and 131-03-9 from 22.11.2021).

## Author Contributions

Conceptualization, methodology, writing—original draft preparation, A.V.S., V.R.N. and A.E.N.; sampling, A.V.S., V.R.N., A.E.N., M.M.D, A.M.B., A.I.R. and E.V.V; data analysis, A.V.S., V.R.N., A.E.N., and M.M.D., writing—review and editing, A.V.S., V.R.N., A.E.N., A.I.R. and E.V.V.; funding acquisition, A.V.S. All authors have read and agreed to the published version of the manuscript.

## References

Abe H (1976) Age determination of Clethrionomys rufocanus bedfordiae (Thomas) (In Japanese with a summary in Eng.). Jpn J Ecol 26:221–227

Akimov IA (2009) Red data book of Ukraine. Animals [In Ukrainian]. Global consulting, Kyiv. ISBN 978-966-97059-0-7

Alibhai S (2009) An X-ray technique for ageing Bank voles (Clethrionomys glareolus) using the first mandibular molar. J Zool 191(3):418–423. 10.1111/j.1469-7998.1980.tb01470.x

Armstrong D, Seddon P (2008) Directions in reintroduction biology. Trends Ecol Evol 23(1):20–25. 10.1016/j.tree.2007.10.003

Barratclough A, Sanz-Requena R, Marti-Bonmati L, Schmitt TL, Jensen E, García-Párraga D (2019) Radiographic assessment of pectoral flipper bone maturation in bottlenose dolphins (Tursiops truncatus), as a novel technique to accurately estimate chronological age. PLoS One 14(9):e0222722. 10.1371/journal.pone.0222722

Brunet-Rossinni AK, Wilkinson GS (2009) Methods for age estimation and the study of senescence in bats. In: Kunz TH, Parsons S (eds) Ecological and behavioral methods for the study of bats. Johns Hopkins University Press: Baltimore, Maryland, pp. 315–325

Buchanan K, Burt de Perera T, Carere C et al (2012) Guidelines for the treatment of animals in behavioural research and teaching. Anim Behav 83:301–309. 10.1016/j.anbehav.2011.10.031

Caspar K, Müller J, Begall S (2021) Effects of sex and breeding status on skull morphology in cooperatively breeding Ansell’s mole-rats and an appraisal of sexual dimorphism in the Bathyergidae. Front Ecol Evol 9. 10.3389/fevo.2021.638754

Caspar KR, Stopka P, Issel D, Katschak KH, Zöllner T, Zupanc S, Žáček P, Begall S (2022) Perioral secretions enable complex social signaling in African mole-rats (genus Fukomys). Sci Rep, 12(1): 22366. 10.1038/s41598-022-26351-3

Cook RD (1979) Influential observations in linear regression. J Am Stat Assoc 74(365):169–174. 10.1080/01621459.1979.10481634

Dunaeva TN (1955) Towards the study of the breeding biology of the common shrew (Sorex araneus L.). Bull Soc Nat Mosc 60 (6):27–43 [in Russian]

Dymskaya MM, Volodin IA, Smorkatcheva AV, Rudyk A, Volodina EV (2024) Field experiments disclose acoustic variation of sonic and ultrasonic calls in a subterranean rodent, the northern mole-vole Ellobius talpinus. J Mammal submitted

Ecke D, Kinney A (1956) American society of mammalogists aging meadow mice, Microtus californicus, by observation of molt progression. J Mammal 37(2): 249–254

Evdokimov NG (1997) Methodology for determining the growth of the mole vole Ellobius talpinus (Rodentia, Cricetidae) Zool J Russ Acad Sci 76(9):1094-1101 [in Russian]

Evdokimov NG (2001) Population ecology of the mole-vole. Yekaterinburg, Yekaterinburg [in Russian].

Flinn EB, Strickland BK, Demarais S, Christiansen D (2013) Age and gender affect epiphyseal closure in white-tailed deer. Southeast Nat 12(2) 297–306. 10.1656/058.012.0205

Gilbert F, Stolt S (1970) Variability in aging Maine white-tailed deer by tooth-wear characteristics. J Wildl Manage 34(3):532–535.

Golov B (1954) Live-trap for the mole vole. Bull Soc Nat Mosc 59(5):95–96 [in Russian]

González-Esteban J, Villate I, Castién E, Rey I, Gosálbez J (2002) Age determination of Galemys pyrenaicus. Acta Theriol 47(1):107–112. 10.1007/BF03193570

Gustafsson TO, Andersson CB, Westlin LM (1982) Determining the age of bank voles - a laboratory study. Acta Theriol 27(20), 275–282.

Hawkins MG, Pascoe PJ (2020) Anesthesia, analgesia, and sedation of small mammals. In: Quesenberry KE, Carpenter JW, Orcutt CJ, Mans C (eds) Ferrets, rabbits and rodents clinical medicine and surgery, 4th ed. Elsevier Saunders, St Louis, MO, USA, pp 536–558. 10.1016/B978-1-4160-6621-7.00031-2

Hecht L (2021) The importance of considering age when quantifying wild animals’ welfare. Biol Rev 96(6):2602–2616. 10.1111/brv.12769

Holmes EE, York AE (2003) Using age structure to detect impacts on threatened populations: a case study with Steller sea lions. Conserv Biol 17(6):1794–1806.10.1111/j.1523-1739.2003.00191.x

Hulejová Sládkovičová H.V, Žiak D, Miklós P, Kameniar O, Kocian Ľ (2019) Age determination and individual growth rate of Microtus oeconomus mehelyi based on live-trapping. Biol 74(5):487–492. 10.2478/s11756-018-00188-6

Jenks J, Bowyer R, Clark A (1984) Sex and age-class determination for Fisher using radiographs of canine teeth. J Wildl Manage 48(2):626–628.

Karels T, Bryant A, Karels D, Bryant T, Hik A, Karels T, Hik D (2004) Comparison of discriminant function and classification tree analyses for age classification of marmots. Oikos 105(3):575–587. 10.1111/j.0030-1299.2004.12732.x

Kaya A, Coşkun Y (2015) Reproduction, postnatal development, and social behavior of Ellobius lutescens Thomas, 1897 (Mammalia: Rodentia) in captivity. Turk J Zool, 39(3), 425–431. 10.3906/zoo-1401-73

Klenova AV, Chelysheva EV, Vasilieva NA, Volodin IA, Volodina EV (2023) Acoustic features of long-distance calls of wild cheetahs (Acinonyx jubatus) are linked to the caller age from newborns to adults. Ethology. 10.1111/eth.13406

Klevezal G (2007) Principles and methods of determination of the age of mammals. KMK, Moscow [in Russian]

Krämer W, Sonnberger H (1989) The linear regression model under test. J Appl É conom 4(2):209–211.

Krebs CJ (1999) Ecological methodology. Addison-Wesle, New York

Kropacheva YE, Cheprakov MI, Sineva NV, Evdokimov NG, Kuzmina EA, Smirnov NG (2018) Dimensions of the body and molars in the mole vole (Ellobius talpinus, Rodentia, Cricetidae) depending on the age and environmental conditions. Biol Bull Russ Acad Sci 45(7):766–771 [in Russian] 10.1134/S1062359018070099

Kropacheva YE, Smirnov NG, Zykov SV (2021) Growth rate of cheek teeth in narrow-skulled vole (Lasiopodomys gregalis) depending on food abrasiveness. Russ J Ecol 52(6):496–503. 10.1134/S1067413621060072

Kryštufek B, Kolarič K, Paunović M (2000) Age determination and age structure in Martino’s vole Dinaromys bogdanovi. Mammalia 64(3):361–370. 10.1515/mamm.2000.64.3.361

Kuprina KV, Smorkatcheva AV (2019) Noninvasive age estimation in rodents by measuring incisors width, with the Zaisan mole vole (Ellobius tancrei) as an example. Mammalia 83(1):64–69. 10.1515/mammalia-2017-0163

Landon DB, Waite CA, Peterson RO, Mech LD (1998) Evaluation of age determination techniques for gray wolves. J Wildl Manage 62(2):674–682.

Letitskaya E (1984) Materials on reproduction and postnatal development of the northern mole vole, Ellobius talpinus (Rodentia, Cricetidae). Zool Zh 63(7):1084–1089.

Lichti N, Kellner KF, Smyser TJ, Johnson SA (2017) Bayesian model-based age classification using small mammal body mass and capture dates. J Mammal 98(5):1379–1388. 10.1093/jmammal/gyx057

Lowe V (1971) Root development of molar teeth in the bank vole (Clethrionomys glareolus). J Anim Ecol 40(1):49–61.

Mazerolle M (2023) AICcmodavg: Model selection and multimodel inference based on (Q)AIC(c). R package version 2.3.-3).

Morris P (1972) A review of mammalian age determination methods. Mammal Rev 2(3):69–104.

Moshkin MP, Novikov E, Petrovski D (2007) Skimping as an adaptive strategy in social fossorial rodents: the mole vole (Ellobius talpinus) as an example. In: Begall S, Burda H, Schleich CE. Subterranean rodents: news from underground (eds). Springer, Heidelberg, Germany. pp. 49–60

Nicholson TE, Mayer KA, Staedler MM, Gagné TO, Murray MJ, Young MA, Tomoleoni JA, Tinker MT, Van Houtan KS (2020) Robust age estimation of southern sea otters from multiple morphometrics. Ecol Evol 10(16):8592–8609. 10.1002/ece3.6493

Olenev G (1989) Functional determination of ontogenetic changes in age markers in rodents and their practical utilization in populational investigations. Sov J Ecol 20:76–86.

R Core Team (2017) R: a language and environment for statistical computing. Version 4.3.1

Read FL, Hohn AA, Lockyer CH (2018) Age estimation methods in marine mammals: with special reference to monodontids. NAMMCO Sci Publ 10. 10.7557/3.4474

Royston P (1995) Remark AS R94: A remark on algorithm AS 181: The W-test for normality. J R Stat Soc C 44(4): 547–551.

Smorkatcheva AV, Kumaitova AR, Kuprina KV (2016) Make haste slowly: Reproduction in the Zaisan mole vole (Ellobius tancrei). Can J Zool 94(3):155–162. 10.1139/cjz-2015-0051

Smorkatcheva AV, Kuprina KV (2018) Does sire replacement trigger plural reproduction in matrifilial groups of a singular breeder, Ellobius tancrei? Mamm Biol 88:144–150. 10.1016/j.mambio.2017.09.005.

Spinage C (1973) A review of the age determination of mammals by means of teeth, with especial reference to Africa. Afr J Ecol 11:165–187.

Stander P (1997) Field age determination of leopards by tooth wear. Afr J Ecol 35:156–161.

Tarasov SA (1966) Determnation of field-vole age by the roentgen technique Zool Zh 45(8): 1247–1250 [in Russian].

Tupikova NV, Sidorova GA, Konovalova EA (1968). A method of age determination in Clethrionomys. Acta Ther 13: 99–115

Viitala J (1971) Age determination in Clethrionomys rufocanus (Sundevall). Ann Zool Fenn 8(1):63–67.

Zadubrovskaya IV, Zadubrovskii PA, Novikov EA (2020) Reproductive characteristics of the northern mole vole at the northeastern periphery of species range. Russ J Ecol 51(2):151–156. 10.1134/S1067413620010142

Zeiler J (1988). Age determination based on epiphyseal fusion in post-cranial bones and tooth wear in otters (Lutra lutra). J Arch Sci 15(15) 555–561. 10.1016/0305-4403(88)90082-9

Zhao M, Klaassen CAJ, Lisovski S, Klaassen M (2019) The adequacy of aging techniques in vertebrates for rapid estimation of population mortality rates from age distributions. Ecol Evol 9(3):1394–1402. 10.1002/ece3.4854

Zudova GA, Proskurnyak LP, Nazarova GG (2017) Age and sex determination in the water vole (Arvicola amphibius, Rodentia, Arvicolinae) based on measurements of pelvic limb bones. Zool J Russ Acad Sci 96:606–613. 10.7868/S0044513417050117

