## Supplementary material for "Dental radiography as a low-invasive field technique to estimate age in small rodents, with the mole voles (*Ellobius*) as an example": Skull radiography of live, non-sedated mole voles

For the manuscript by Nikonova et al.

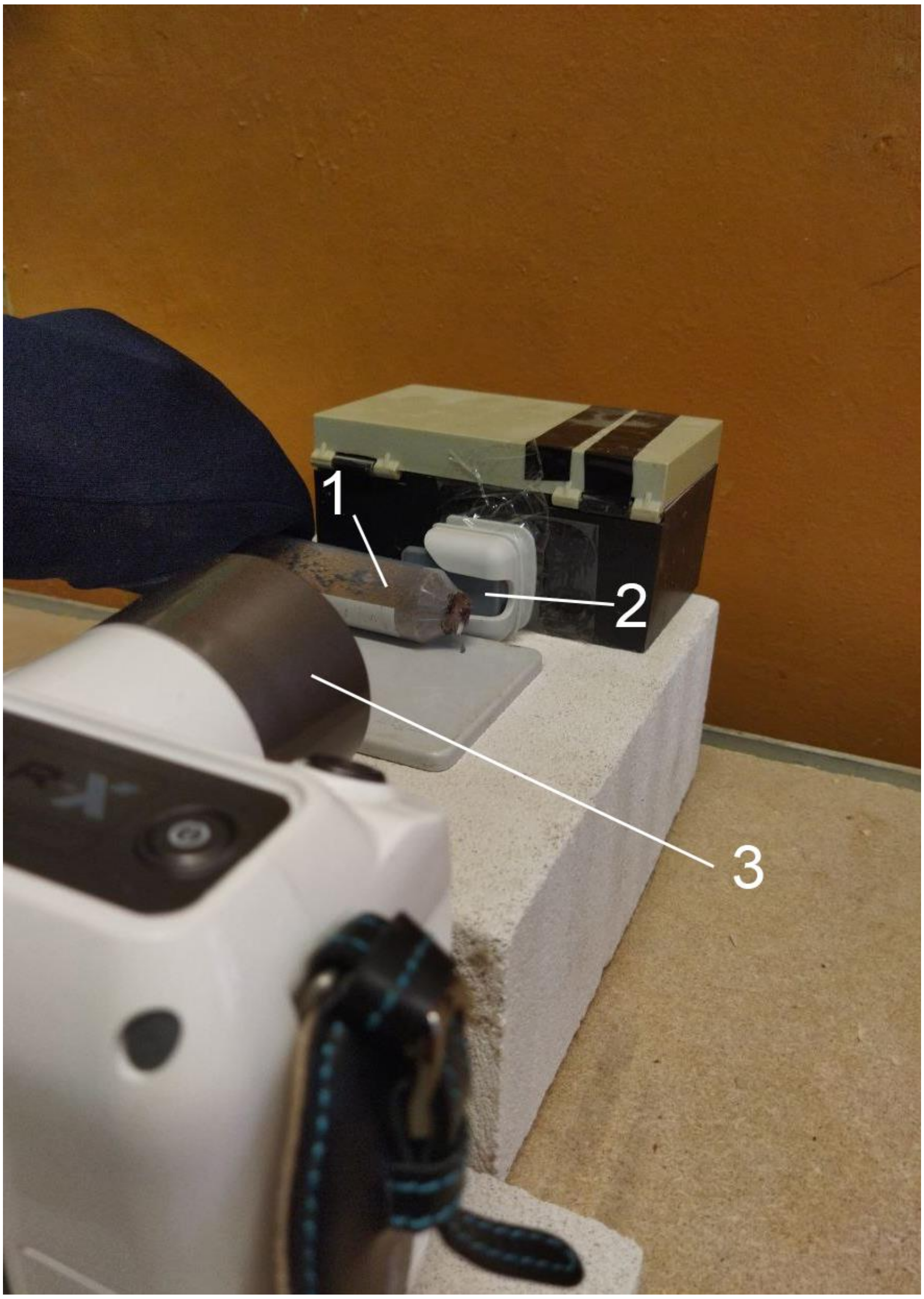


Fig. SI1. Skull radiography of a live, non-sedated mole vole.

1 - falcon tube with an animal
2 - dental Radiovisiography Sensor EzSensor 1.5 (Vatech);
3 - portable X-ray equipment (Rexstar LCD, Korea).

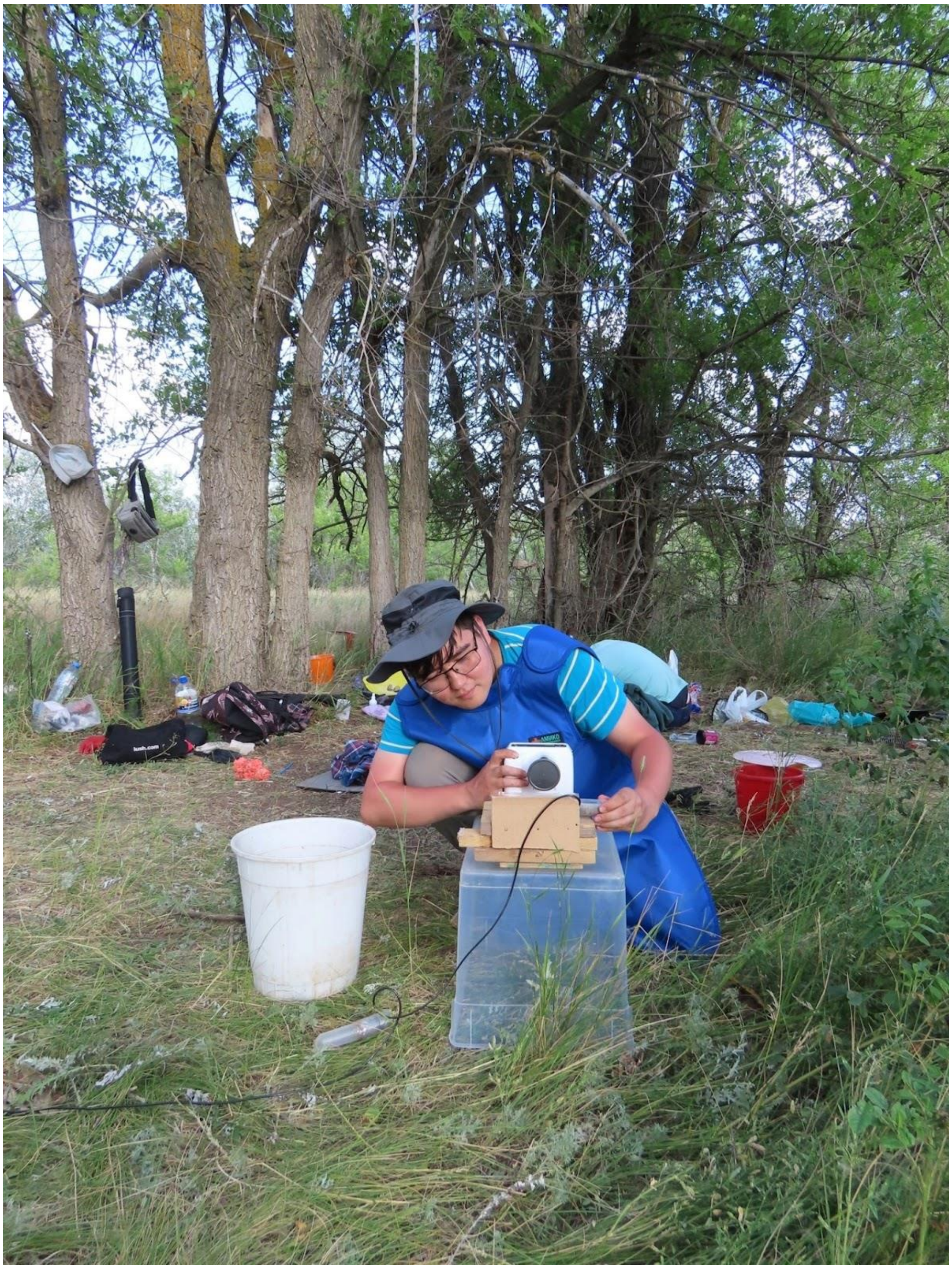

Fig. SI2. Radiography of a wild mole vole in field
